# Adrenocortical Cancer Cell uptake of Iron Oxide Nanoparticles

**DOI:** 10.1101/2024.12.04.626790

**Authors:** Ritihaas Surya Challapalli, Cong Hong, Anna Sorushanova, Obdulia Covarrubias-Zambrano, Nathan Mullen, Sarah Feely, Jose Covarrubias, Sunita N. Varghese, Constanze Hantel, Peter Owens, Martin O’Halloran, Punit Prakash, Stefan H. Bossmann, Michael Conall Dennedy

**Affiliations:** Discipline of Pharmacology & Therapeutics, School of Medicine, University of Galway, Galway, Ireland; Department of Cancer Biology, The University of Kansas Medical Center, Kansas City, KS. USA; Department of Endocrinology, Diabetes, and Clinical Nutrition, University Hospital Zurich, Zurich, Switzerland; Medizinische Klinik und Poliklinik III, University Hopsital Carl Gustav Carus Dresden, Dresden, Germany; Centre for Microscopy & Imaging, University of Galway, Ireland; Translational Medical Device Lab, University of Galway, Galway, Ireland; Department of Electrical and Computer Engineering, Kansas State University, Manhattan, KS. USA; Department of Biomedical Engineering, The George Washington University, Washington, DC, USA

**Keywords:** Nanoparticles, HAC15, MUC-1, H295R, HUVEC, iron oxide, monocytes

## Abstract

Adrenocortical carcinoma (ACC) is a rare cancer with poor prognosis, treated primarily through surgery and chemotherapy. Other treatments like radiation or thermal ablation for metastases have limited success, and recurrence is common. More effective management options are needed. Magnetic iron oxide nanoparticles (IONP) show promise in cancer treatment due to their ability to be modified for selective uptake by cancer cells. This study investigated IONP uptake in ACC cell lines (H295R, HAC-15, MUC-1) using a multicellular model with endothelial cells (HUVEC) and monocytes. IONP uptake was concentration- and time-dependent, with optimal uptake at 10 µg/mL. IONP were found in the cytoplasm and intracellular vesicles of ACC cells. However, endothelial cells and monocytes also absorbed IONP, reducing uptake by ACC cells. These findings suggest ACC cells actively take up IONP, but better targeting is needed to enhance uptake specificity and efficiency.

## Introduction

Adrenocortical carcinoma (ACC) is a rare malignancy which carries a poor prognosis (median survival 15-24 months) and limited treatment options. Five-year survival for patients with ENS@T Stage III disease or above is <20% [1]. For those with Stages I and II disease R0 surgical resection offers the potential for cure. However, recurrence is high and in those who recur, or have higher disease stages, mitotane and combination chemotherapy have limited success [1,2]. Efficacy of radiotherapy is debated while local ablation of individual metastases for disease control has been met with limited success [3,4]. Mitotane is the primary licenced chemotherapy for adjuvant management [5] and is also part of commonly used combination chemotherapy regimens, namely Mitotane/Etoposide/Cisplatin/Doxorubicin (M-EDP) [5,6] and Mitotane-Streptotozocin [7]. However, mitotane has a narrow therapeutic window (14-20mg/L) and even at therapeutic concentrations, tolerability is poor and toxicity high [5]. Therefore, improved therapeutic options for ACC are needed, whether this takes the form of better delivery of existing cytotoxic regimens to achieve higher cellular concentrations or the development of advanced combination cytotoxic strategies to combat disease. One method for improving cytotoxic efficacy when treating cancers is nanoparticle-mediated, e.g. targeted delivery of chemotherapy or thermal-activated cytotoxic therapy [8].

Nanoparticles offer potential as diagnostic and therapeutic tools (so-called theranostic agents) in the management of cancer[8]. Functionalised nanoparticles have been used in applications such as drug delivery, cancer immunotherapy and diagnosis [9], delivery of hyperthermia [10], bioimaging, cell labelling and gene delivery [8,11,12]. Nanotechnology can be engineered in various shapes and sizes, and has the capability for modification of surface characteristics using ligand-coating of molecules, which can selectively target individual cell types for selective uptake [8]. These properties can be exploited for cancer chemotherapy to embed cytotoxic agents within the nanoparticle core and deliver them to individual tumour cells at much higher concentrations than traditional systemic chemotherapy [8,9]. Once at the tumour site, nanoparticles can be stimulated to release their contents immediately or in a controlled manner, minimising systemic adverse effects.

Magnetic metal nanoparticles have additional utility as nanobiocontrast agents, which can be imaged using CT and MRI [8,10,12]. In turn, these agents offer the advantage of dual theranostic properties whereby their concentration within the target tumours can be imaged and monitored. Magnetic metal nanoparticles can also be activated to release their contents or stimulated to deliver other cytotoxic or adjuvant therapies such as hyperthermia, using light or magnetic resonance, respectively [8,10].

While nanoparticles have the potential to be delivered specifically to cancer cells and thereby deliver targeted therapy, the overall bio-interaction of these cells within a complex multicellular environment must also be considered. Nanoparticles are typically administered systemicallyand travel through the circulation to their intended target tissue [13] or locally via intra-arterial administration [14]. Therefore, before reaching their target, these molecules will have interacted with cellular blood components, the endothelium and connective tissue components [13]. When understanding nanoparticle uptake in the context of cancer, it is also critical to evaluate this in the context of uptake into cellular blood components e.g. nanoparticle interactions with cellular blood components.

The uptake and effects of iron oxide nanoparticle (IONP) uptake into ACC cells has not been evaluated in detail. In the current study, we have evaluated the uptake of IONP into three ACC cell lines, H295R, MUC-1 and HAC15 cells. We investigated cellular nanoparticle uptake, toxicity and effects on cellular function/steroidogenesis. We have also compared uptake of IONP in the presence and absence of endothelial cells and human primary monocytes.

## Materials and methods

### Materials

HAC15 and H295R cells were purchased from ATCC. HUVEC cells were purchased from Lonza. MUC-1 cells were provided by Dr. Constanze Hantel, University of Zürich, Switzerland [15]. Nu serum and ITS+ were purchased from Corning. Cosmic calf serum was purchased from Fisher Scientific.

### Adrenocortical carcinoma cell culture

Human adrenocortical carcinoma (ACC) cells (MUC-1, H295R and HAC15) were seeded in 24 well plates and media were changed every 2 days. MUC-1 cells were cultured for 3 days in Advanced DMEM: F12, supplemented with 10 % FBS and 1 % penicillin/ streptomycin at 37 °C and 5 % CO_2_. H295R cells were cultured for 3 days in DMEM: F12, supplemented with 2.5 % Nu serum, 1 % ITS+ and 1 % penicillin/ streptomycin at 37 °C and 5 % CO_2_. HAC15 cells were cultured for 3 days in DMEM: F12, supplemented with 10 % CCS, 1 % ITS+ and 1 % penicillin/ streptomycin at 37 °C and 5 % CO_2_.

### Endothelial cell culture

Human endothelial cells (HUVEC) were seeded in 24 well plates and media were changed every 2 days. Cells were cultured for 3 days in Endothelial media with ready-to-use supplement mix (fetal calf serum, endothelial cell growth supplement, epidermal growth factor, heparin and hydrocortisone) at 37 °C and 5 % CO_2_ prior to addition of Magnetic Iron Oxide Nanoparticles (IONP).

### Primary monocyte isolation and Cell Culture

Peripheral blood was obtained from adult donors (aged 18-68 years) at the hemochromatosis clinic during venesection at University Hospital Galway, with prior informed consent from all donors in accordance with the University of Galway Research Ethics Committee. All donors had a diagnosis of uncomplicated hemochromatosis but had normal iron studies and were undergoing prophylactic venesection. Patients were otherwise healthy and without co-morbidity.

Peripheral blood mononuclear cells (PBMCs) were isolated using Ficoll®-Paque Premium (Sigma Aldrich) gradient centrifugation method from blood of healthy donors. Untouched monocytes were isolated from PBMCs using Pan Monocyte Isolation Kit and an LS Column (Miltenyi Biotec). Isolated Monocytes were then characterised for surface marker expression using PerCP-Cy™5.5 Mouse Anti-Human CD14 Clone MΦP9 (RUO), BV786 Mouse Anti-Human CD16 Clone 3G8 (RUO), FITC Mouse Anti-Human CD45 Clone HI30 (RUO) and BV605 Mouse Anti-Human HLA-DR Clone G46-6 (RUO) (BD Biosciences) on Cytek® Northern Lights™ 2000 Flow Cytometer. Sort efficiency and purity was routinely validated using Cytek® Northern Lights™ 2000.

### Fe/Fe_3_O_4_ core/shell Nanoparticles

Magnetic Fe/Fe_3_O_4_ core/shell nanoparticles were synthesized at the University of Kansas Medical Center in accordance with a published procedure[16]. In short, Iron pentacarbonyl Fe(CO)_5_ was thermally decomposed in the presence of oleylamine and hexadecylammonium chloride (HADxHCl) using 1-octadecene (ODE) as solvent. After controlled air oxidation and exchange of surface-bound oleylamine vs. dopamine, nanoparticles with a well-defined core/shell structure (average Fe(0) core diameter of 12+/-1.5 nm and the Fe_3_O_4_ shell thickness of 3.0+/-1.5) were obtained. Dopamine forms stable organic coatings with binding constants of the order of 10^15^ l mol^−1^ on Fe_3_O_4_ [17]. High resolution -TEM (HRTEM) reveals the polycrystalline nature of the nanoparticles.

### Peptide Synthesis

K_20_G was synthesized via standard Solid Phase Peptide Synthesis (SPPS) [18]. Briefly, preloaded trityl-resin was swelled in DCM for 20 min, after washing with DMF, Fmoc-protected amino acids were added sequentially with O-Benzotriazole-N,N,N’,N’-tetramethyl-uronium-hexafluoro-phosphate (HBTU) as coupling agent in a mixture of diisopropylethylamine (DIEA) and DMF. The purity of the peptide was ascertained to be > 95% by means of HPLC/MS (quadrupole).

### Assembly of the IONPs

200 mg of dopamine coated Fe/Fe_3_O_4_ nanoparticles were dispersed in 5.0 mL of DMF (dimethylformamide). To this dispersion, 2 mmol of K_20_G, 2.2 mmol of EDC (1-Ethyl-3-(3-dimethylaminopropyl) carbodiimide), 1 mmol of DMAP (4-(dimethylamino)pyridine) in 2.0 mL of DMF were added. After sonicating for 1 h, the nanoparticles were centrifuged (3000 RPM) and thoroughly washed with DMF (1 mL × 10) and then diethyl ether (1mL x 5). Characterization was performed by means of TEM (Figure 1) and dynamic light scattering (DLS): Dopamine coated Fe/Fe_3_O_4_: hydrodynamic diameter 209 ± 1.5 nm, polydispersity: 0.091; K20G-dopamine coated Fe/Fe_3_O_4_: hydrodynamic diameter 383 ± 18 nm, polydispersity: 0.293.

**Figure 1:**
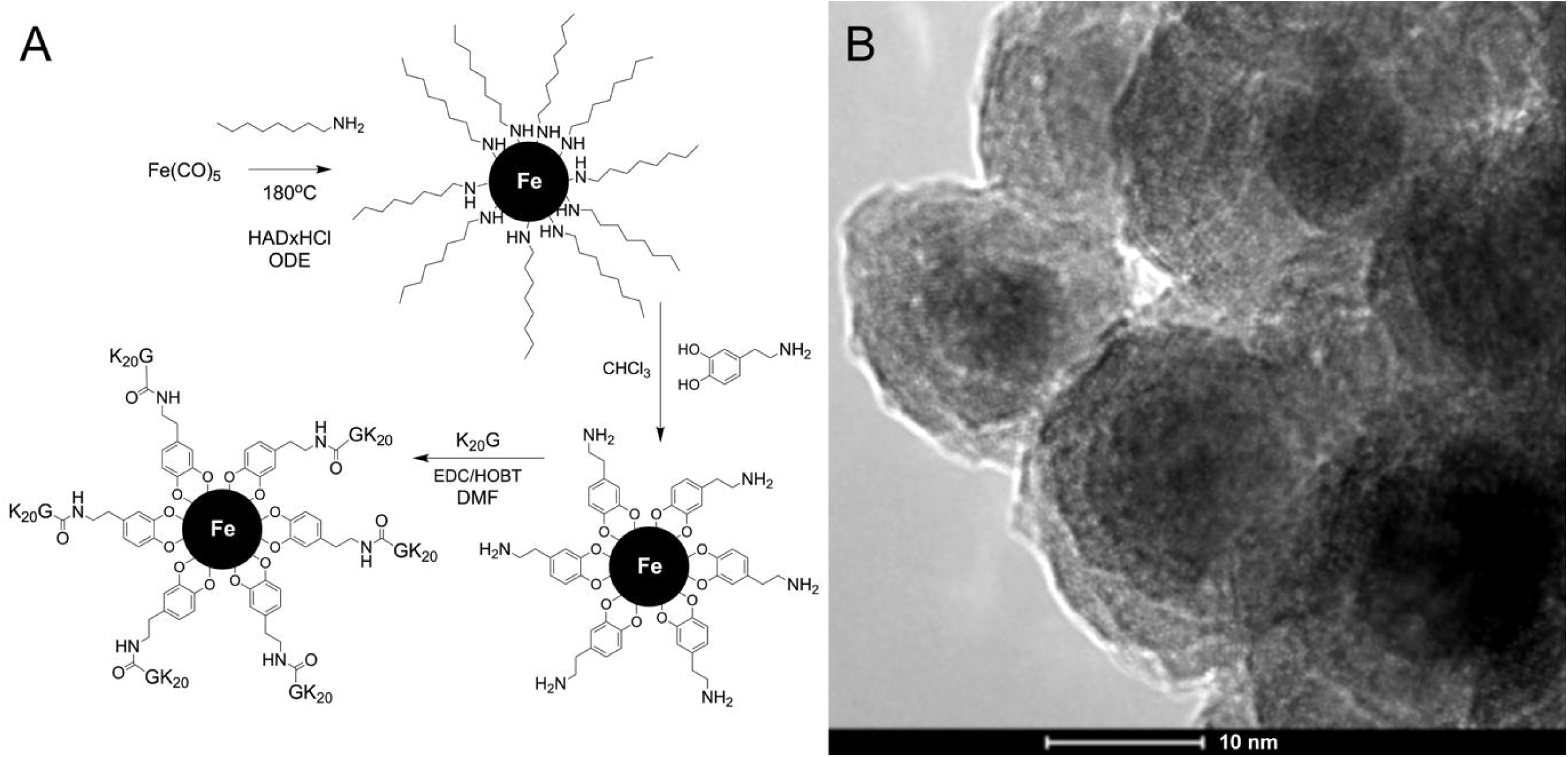
(A) Synthesis scheme and B) TEM images of assembled IONP.

### Transwell Cell Culture System

A transwell cell culture system (Greiner ThinCert cell culture inserts, pore size: 8.0 µm) was used to assess IONP uptake by the ACC cells in the presence and absence of primary monocytes with a fibronectin, and a fibronectin and HUVEC layer. ACC cells were seeded at the bottom of the well and cultured for 3 days. Transwell migration inserts were coated with fibronectin overnight. HUVEC cells were seeded on the fibronectin coated insert and cultured for 3 days. Briefly, the inserts were added into the wells with ACC cells with fresh media after 3 days in culture. Primary monocytes and IONP at concentration of 10 µg/ml were added to the top of the insert and incubated for 24 hours. Flow cytometry was used to evaluate IONP uptake by the primary monocytes in the supernatant at the top of the insert and at the bottom of the well, HUVEC layer on the insert and the ACC cells at the bottom of the well.

### Viability, nanoparticle uptake and rate of uptake by ACC, endothelial cells and Primary monocytes

Flow cytometry (Cytek, Northern Lights 2000®) and the associated acquisition software (SpectroFlo® Version 3.0.3) was used to: (i) determine cellular viability and (ii) IONP uptake by ACC cell lines, HUVEC endothelial cells and primary monocytes. ACC cells and HUVEC Endothelial cells were treated with IONP suspended in fresh cell culture media at concentrations of 0.5, 5, 10, 20 and 50 µg/ml, and incubated for 24 hours. After 24 hours of incubation, cells were washed with DPBS and trypsinised (0.25% trypsin-EDTA). Primary Monocytes were treated with IONP concentrations of 5, 10, 20 µg/ml for 24 hrs. IONP were tagged with rhodamine and uptake by live cells was analysed using flow cytometry. Cellular viability was determined using 10 µM Sytox blue staining of each cell type.

Median fluorescence intensity (MFI) was used to determine the loading of the IONP in the cells compared to the control. All experiments were performed in triplicate unless otherwise specified.

For *rate-of-uptake experiments*, Optimum concentration was determined at 10 µg/mL and was used to evaluate IONP uptake rate by ACC cells. IONP were added to the cells at 1-6 hour time-points to determine the rate of uptake and cells were analysed by flow cytometry. Data was analysed by using FCS Express (De Novo Software) and FlowJo® (v 9.0 Treestar, BD Biosciences).

The residual uptake of IONP (10µg/mL) and its effect on cellular viability across all three ACC cell lines, HUVEC and primary human monocytes was also investigated following 7 days in culture. Cells were first exposed to IONP in cell culture media for 24 hours, following which media were changed. Media was then changed every 2 days thereafter, and the cells were analysed by flow cytometry as described above.

### Ki-67 expression of ACC and endothelial cells

Ki-67 assay was performed to assess cellular proliferation using PE anti-mouse/Human Ki-67 (Biolegend). Briefly, supernatants were removed post-treatment, and cells were washed with PBS and trypsinised. 100,000 cells were isolated and centrifuged for 5 minutes at 400 RCF. Cells were fixed in 4% formalin at room temperature for 20 minutes. Cells were then washed with FACS buffer, spun down and resuspended in Intracellular staining buffer. Ki-67 stain was made up in FACS buffer and cells were resuspended and stained for 15 minutes in the dark at 4 C°. Following washing steps, cells were analysed by flow cytometry (Cytek, NL 2000®).

Analysis was carried using on FCS Express® (De Novo Software) and FlowJo® (v 9.0, Treestar, BD Biosciences)

### Metabolic and mitochondrial activity of ACC and endothelial cells

Cellular metabolic activity was evaluated using the Alamar Blue assay. Cells were seeded and media was removed on day 3 of culture and IONP were added in fresh media to the cells at concentrations of 0.5, 5, 10, 20 and 50 µg/ml, and incubated for 24 hours. Post incubation, 10% of Alamar Blue was added to the cells and incubated at 37°C for 3 hours. Fluorescence was read at excitation wavelength of 560 nm and an emission wavelength of 590 nm.

Oxygen consumption rate (OCR), ATP production, non-mitochondrial OC, maximal respiration, spare respiration and basal were measured with a XF24 extracellular analyser (Seahorse Bioscience) and XF Cell Mito Stress Test Kit (Seahorse Bioscience). Cells were seeded and cultured in XF24 plates in DMEM media supplemented with 10% FBS, 25mM glucose, 2mM glutamine for 24 hours prior to incubation with IONP at concentration of 10 µg/ml for further 24 hours. One hour prior to the experiment, 1 ml of XF calibrator was added to each well of the XF cartridge and incubated at 37 °C and 0% CO2. Cells were washed with PBS and respective XF assay media was added per well and incubated for 1 hours at 37 °C and 0% CO2. The hydrated cartridge and the plate were removed from the incubator. Oligomycin and FCCP were added to the injection ports of the XF cartridge. Experiment was initiated through the instrument interface.

### Microscopy and TEM of ACC, endothelial cells and primary monocytes

IONP uptake in ACCs, endothelial cells and primary monocytes was visualised via confocal microscopy (FV3000, Olympus Fluoview Laser scanning confocal microscope, incubation at 37C, 5% CO_2_ control) following 24 hours of incubation. ACCs and endothelial cells were incubated with IONP at concentrations of 0.5, 5, 10, 20 and 50 µg/ml for 24 hours, while primary monocytes were incubated with 5, 10 and 20 µg/ml for 24 hours. Media containing IONP was removed after 24 hours of incubation and the cells were washed with DPBS and fixed with 4 % paraformaldehyde (PFA). To visualise IONP rate of uptake, cells were incubated with optimum IONP concentration of 10 µg/ml for 1-6 hours. Media containing IONP was removed after incubation, the cells were washed with DPBS and fixed with 4 % paraformaldehyde (PFA).

Cellular viability was assessed following 24-hour incubation with IONP at specified doses. Following incubation, the cells were stained with calcein AM (live) and ethidium homodimer-1 (dead) for 30 minutes and images were acquired at random locations of well using EVOS® Cell Imaging System M7000 (Thermo Scientific).

For cell morphology assessment, media was removed after 24 hours of incubation with IONP and the cells were washed with DPBS. Cells were fixed with 4 % paraformaldehyde (PFA), permeabilised with 0.2 % Triton X-100 and then nuclei were stained with 4′,6-diamidino-2-phenylindole (DAPI) and cytoskeleton was stained with Rhodamine FITC-488. Morphology was assessed using EVOS® Cell Imaging System M7000 (Thermo Scientific).

To determine intracellular location of IONP, cells were incubated with IONP at concentration of 10 µg/ml for 24 hours. Cells were washed with DPBS and fixed with EM fixative (2% glutaraldehyde + 2% paraformaldehyde in 0.1M sodium cacodylate buffer pH 7.2) for 2 hours at room temperature. Following fixation, cells were kept in EM buffer (0.2 M sodium cacodylate buffer pH 7.2) at 4 °C until processing. Cells were scraped and the pellet was transferred to Eppendorf tubes. Secondary fixation (1% osmium tetroxide in 0.1M sodium cacodylate buffer pH 7.2) was carried out for 2 hours at room temperature. Cells were dehydrated through a graded series of ethanol (30%, 50%, 70%, 90% and 100%) for 15 minutes, repeated twice. Following final ethanol dehydration step, acetone was added to the cells for 20 minutes, repeated twice. Cells were then infiltrated with 50:50 and 75:25 resin: acetone mixtures for 4 hours, and 100% resin for 6 hours. Cell samples were then polymerised in the oven at 65 °C for 48 hours prior to sectioning and TEM was performed using Hitachi H7500.

ACC, primary monocytes and endothelial cells were treated with IONP at a concentration of 10 µg/ml in an environmental chamber at 37°C and 5% CO_2_ and live cell imaging was performed every 15 mins at three random locations per well over 24hrs using confocal microscopy.

Optimum IONP concentration of 10 µg/ml was used to evaluate the IONP uptake and intracellular location following 7 days in culture. Cells were incubated with IONP for 24 hours and media was changed every 2 days. To visualise IONP uptake, media was removed after 24 hours of incubation with IONP and the cells were washed with DPBS and fixed with 4 % paraformaldehyde (PFA). To visualise intracellular location, TEM was used as described above.

### Steroidogenesis of ACC cells

Quantification of aldosterone and cortisol was performed using HPLC tandem mass spectrometry (SCIEX Q-Trap 4500 liquid chromatography tandem mass spectrometry) operated in MRM mode to determine the effect of IONP on steroid secretion of in H295R and HAC15 cells. Supernatants were extracted from cell culture following treatment, centrifuged in order to remove any possible cell debris, transferred to a fresh tube and stored at -80°C for storage until ready for analysis. The cell number of each sample was calculated for normalization. The interfaced chromatography system was an Agilent 1260 infinity binary pumping with an integrated autoinjector system with Peltier cooled sample storage trays. The column used was a ZORBAX SB-C18 rapid resolution high throughput with the following properties: 2.1 mm (column diameter), 50 mm (column length), 1.8-μm-diameter particles, and kept constantly at 40°C by a temperature-controlled oven. Solvent A was Millipore grade water and solvent B was liquid chromatography mass spectrometry grade acetonitrile, both containing 1% (v/v) mass spectrometer grade formic acid. The HPLC flow rate was 200 μL/minute. The 20-minute HPLC gradient program, along with MRM transition settings, and optimized electrospray and ion settings for cortisol and aldosterone. On the day of the assay, 500 μL of each sample was slowly thawed at 4°C, and then transferred to a 1.5-mL tube. A known concentration of internal standard (500 pg of D4 aldosterone—qmx; ISS11214/100UG/1, 5 ng of D4 cortisol—CDN Isotopes; D-5820) was added to each sample and following a period of equilibration, steroids were extracted using the solvent ethyl acetate (Fisher scientific 15 654 750). After drying, by vacuum enabled centrifugal evaporation, the samples were subsequently reconstituted in 25 μL of acetonitrile and kept at 4°C until ready to run the assay. A 10-point standard curve was constructed using a 1 in 4 dilution for both aldosterone (range: 0.0000572205-15 ng) and cortisol (range: 0.000572205-150 ng). Again, a known concentration of internal standard was added to each sample of the standard curve (500 pg of D4 aldosterone, 5 ng of D4 cortisol) and solid phase extraction was carried out, where the standards were finally reconstituted in 25 μL of acetonitrile prior to acquisition through the HPLC mass spectrometer. The calculated concentrations of both aldosterone and cortisol were measured by assessing the ratio of the detector signal for the analyte and the internal standard of each sample. Limit of detection and limit of quantitation were determined by observing a signal to noise ratio of ≥10:1 or 3:1, respectively. Based on these definitions, for cortisol (analysed in positive ion mode), limit of detection (LOD) and limit of quantitation (LOQ) were estimated to be ∼3 pmol/L and ∼50 pmol/L respectively. For aldosterone (analysed in negative ion mode) LOD and LOQ were estimated to be ∼20 pmol/L and ∼80 pmol/L respectively

### Statistical analysis

The following staining index was used for analysis:

The Resolution Metric: MFI experimental sample - MFI control sample / rSD experimental sample + rSD control sample.

The Staining Metric was used to compare within sample groups or when populations were not normally distributed and calculated as follows: MFI experimental sample - MFI control sample / 2SD control sample

Statistical analyses were performed using GraphPad Prism 10.2 (GraphPad Software, San Diego, CA, USA). Data are represented as mean ± SD unless stated otherwise. Paired sample analyses were performed using a 2-sided Student’s t-test. Multiple-group comparisons were carried out using an unpaired t-test or an analysis of variance (ANOVA) followed by suitable post hoc test, either Dunnett’s or Tukey’s. Statistical significance for 2-tailed analyses (P value) was assigned for values p<0.05.

## Results

### Iron oxide nanoparticle (IONP) are taken up by ACC cells in a concentration and time dependent manner and primarily localised to the cytoplasm

Uptake of the magnetic iron oxide nanoparticles and the rate of uptake of the nanoparticles by ACC cells (MUC-1, H295R, HAC15) at different concentrations was determined by flow cytometry and confocal microscopy. Flow cytometry evaluation of median fluorescence intensity (MFI) showed that the uptake of IONP was concentration-dependent following 24 hrs of incubation. MUC-1 cells showed a higher uptake rate at 24hrs after induction compared to H295R and HAC15 (Figure 2A). Confocal microscopy images revealed that IONP aggregates formed at higher concentrations between 20 and 50 µg/ml (Figure 2C). Considering the potential requirement for intra-arterial localized administration of these agents, aggregates pose the risk of embolus formation and microinfarction. Therefore 10 µg/ml was chosen as the maximal appropriate concentration for theranostically relevant experimentation. IONP at 10 µg/ml also represented the concentration where 50% maximal uptake was observed. Measurement of IONP uptake over 1hr, 2 hrs, 3hrs, 4hrs, 5hrs, 6hrs and 24hrs (Figure 2B) demonstrated time-dependent uptake by all three ACC cell types. IONP were rapidly taken up by MUC-1 cells, with significance demonstrated at 2 hours and increasing intracellular concentrations seen across all timepoints. HAC15 and H295R cells took up IONP more slowly, with significance demonstrated only at 24 hours (Figure 2B). “Real-time” confocal capture (15 minutes intervals over 24 hrs) of IONP uptake (10 µg/ml) by ACC cells validated these findings with increasing fluorescence for each cell type over 24 hrs (Figure 2D).

**Figure 2:**
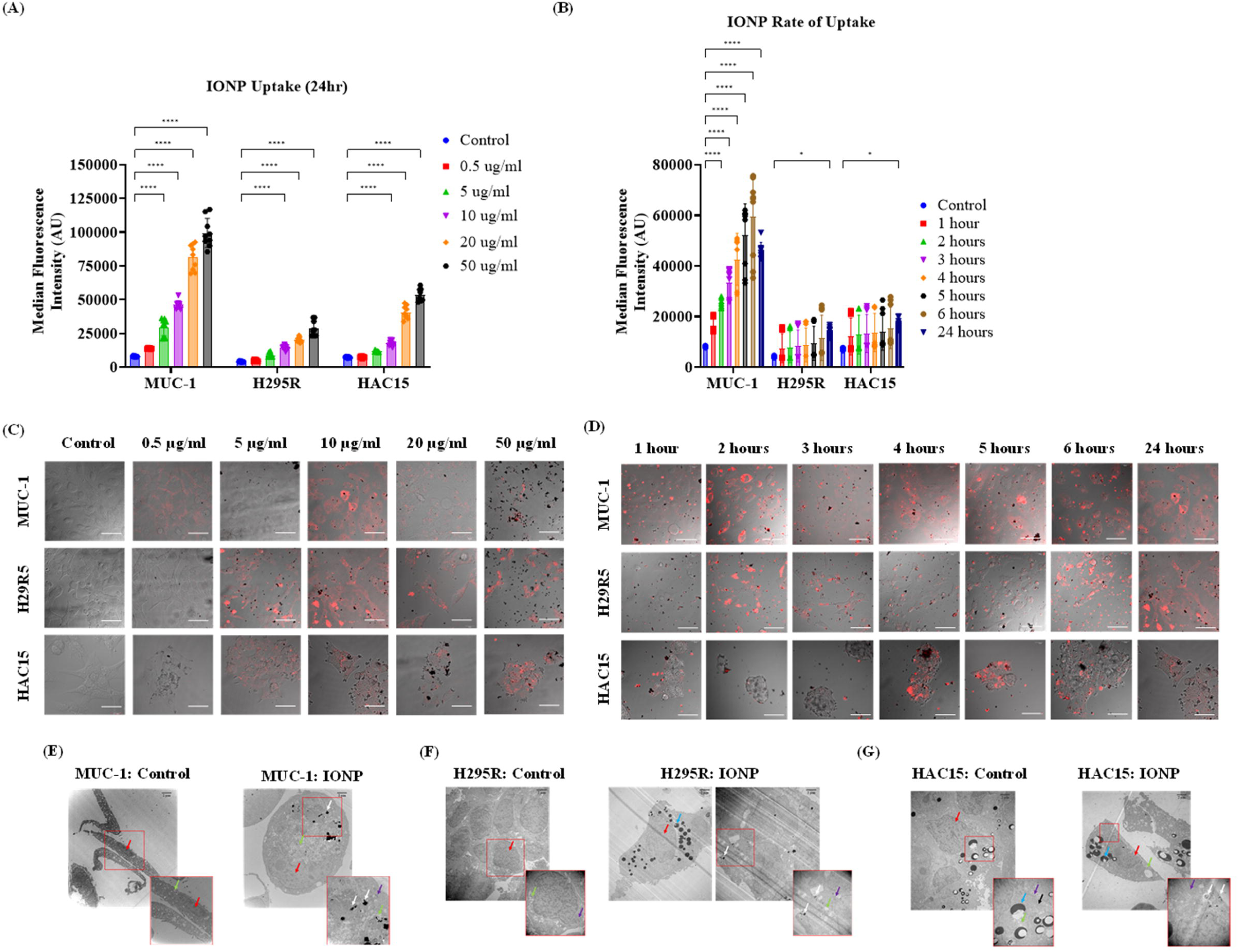
(A) ACC cells were incubated with 0.5, 5, 10, 20 and 50 µg/ml Iron oxide nanoparticle (IONP) uptake was assessed via Flow Cytometry following 24 hrs of incubation; (B) ACC cells were incubated with 10 ug/ml of IONP and rate of uptake was assessed from 1-6 hrs via Flow Cytometry; (C, D) Fluorescent micrographs with brightfield overlay showing Iron oxide nanoparticle (IONP) uptake after 24hrs at different concentrations and rate of uptake following 1-6 hrs and 24 hrs of incubation, respectively, Scalebar: 50 µM. (E-G) Intracellular location of IONPs following incubation with 10 µg/ml of IONP for 24 hrs visualised by Transmission electron microscopy (TEM). White arrow indicates IONP; red arrow indicates nuclei; blue arrow indicates lipids; purple arrow indicates mitochondria; black arrow indicates RER; green arrow indicates vesicles, Scalebar: 2 µM. Data are represented as mean ± SD, (n=3), statistical comparisons were performed using ANOVA analysis, post-hoc. Statistical significance is denoted as *p<0.05, **p<0.01, ***p<0.005,****p<0.001.

TEM was used to visualise the intracellular location of the IONP in each ACC cell type at 24h. In all cell types, TEM images showed that IONP within the cytoplasm or within enzymatic vesicles (Figure 2 E-G) but not in nuclei or mitochondria. Higher amounts of IONP were visible in the MUC-1 (Figure 2E) cells compared to other ACC cell types. Live imaging video capture of IONP uptake over 24 hours demonstrated that ACC cells engulfed IONP by micropinocytosis (Figure S1).

### Iron oxide nanoparticles (IONPs) reduced cell viability and altered morphology at the highest concentration but did not impair ACC cell functionality

The effect of IONP uptake on cellular viability was next assessed by calcein AM, ethidium homodimer-1, using confocal microscopy and Sytox blue using flow cytometry. At IONP concentrations of 50 µg/ml cell viability decreased significantly in both H295R (85.5 ± 10, % p<0.01) and HAC15 (86.3 ±7 % p<0.01) compared with untreated control (Figure 3A). MUC-1 viability interestingly was not affected by IONP at any concentration. Confocal microscopy confirmed these findings with a visible increase of non-viable cells 50 µg/ml IONP for H295R and HAC15 cells but not for MUC-1 (Figure S3).

**Figure 3:**
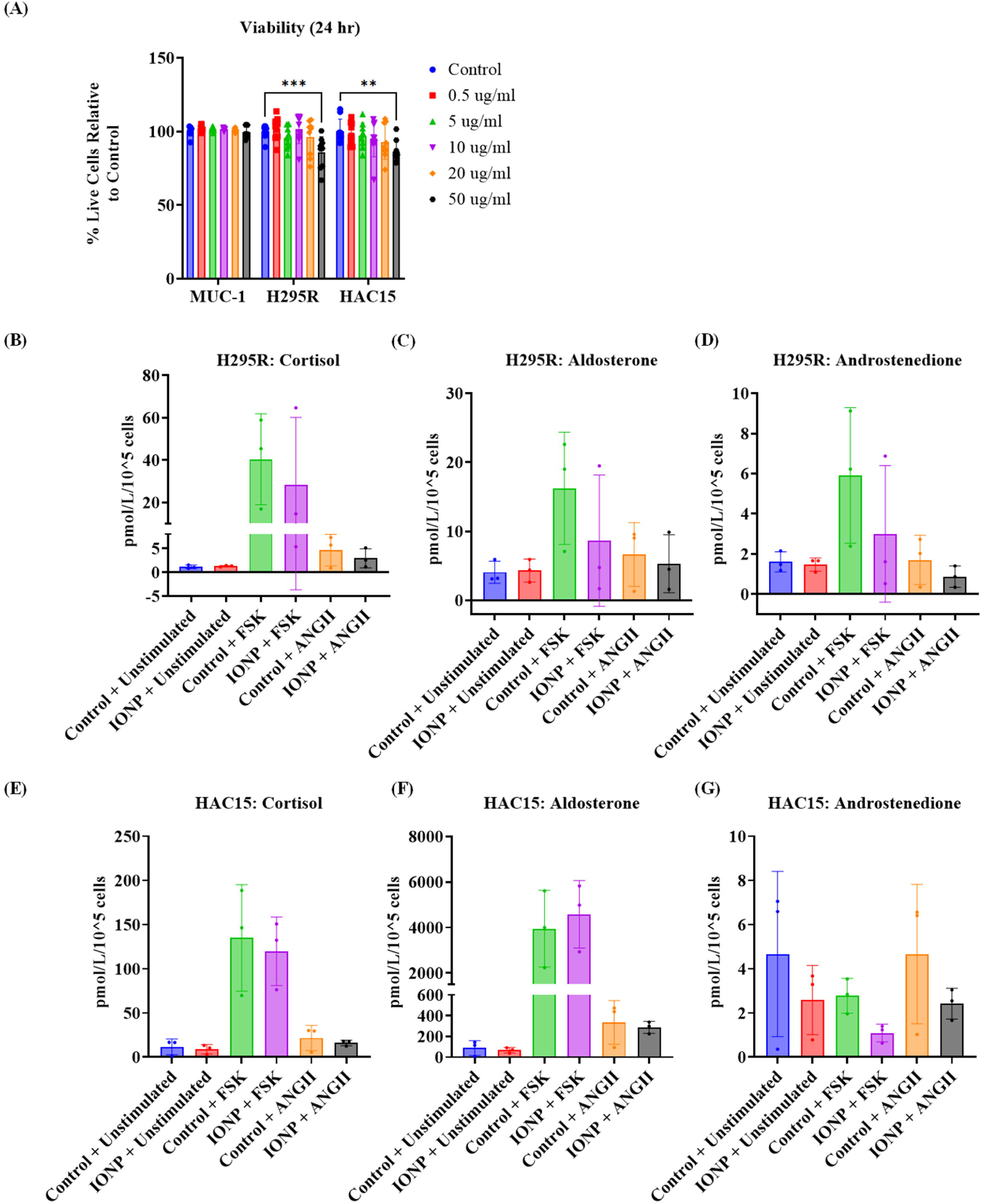
(A) Flow Cytometry analysis of cell viability following incubation with 10 µg/ml of IONP for 24 hrs, expressed relative to untreated control; (B-D) Cortisol, Aldosterone and Androstenedione secretion by H295R following exposure to 10 µg/ml IONP when stimulated with 10 µM Forskolin (FSK) or 10 nM Angiotensin II (Ang II); (E-G) Cortisol, Aldosterone and Androstenedione secretion by HAC15 following exposure to 10 µg/ml IONP when stimulated with 10 µM Forskolin (Fsk) or 10 nM Angiotensin II (Ang II). Data are represented as mean ± SD, (n=3), statistical comparisons were performed using ANOVA analysis, post-hoc. Statistical significance is denoted as *p<0.05, **p<0.01, ***p<0.005, ****p<0.001

Iron excess, e.g. haemochromatosis, in endocrine organs, is associated with the inhibition of hormone synthesis. Therefore, we next assessed the effects of IONP on stimulated adrenal steroidogenesis using LC-MS/MS. Following exposure to 10 µg/ml IONP, forskolin (FSK)-stimulated (10 µM) cortisol and androstenedione secretion, and Angiotensin II (AngII) (10nM)-stimulated aldosterone secretion was measured in H295R and HAC15 cells. At 10 µg/ml IONP there was no effect on steroidogenesis in either cell line (Figure 3 B-G).

Morphological analysis, proliferation and metabolic activity were also examined across all cell types in response to IONP uptake at all concentrations. These data are represented in the supplementary material. In brief, 50 µg/ml affected the morphology (Figure S4) and metabolic activity (Figure S2) of the cells, there was no effect at 10 µg/ml. Proliferation, measured using Ki67, was not affected in H295R or HAC15 cells at any IONP concentration in HAC15 or H295R cells.

### Iron oxide nanoparticle (IONP) are taken up by endothelial cells and primary monocytes

For in vivo use of IONPs as theranostic nanobiocontrast agents, they must be administered intravenously systemically or locally to the artery or adrenal arteries, requiring them to interact with the reticuloendothelial system. Therefore, we next assessed IONP uptake by endothelial cells (HUVECs) and monocytes in the presence and absence of each of the ACC cell lines.

Flow cytometry evaluation of median fluorescence intensity (MFI) of rhodamine-labelled IONP demonstrated a concentration-dependent uptake in both endothelial cells (Figure 4A) and monocytes (Figure 4B) with significance at concentrations of 10 µg/ml and above at 24 h. (Figure 4 A & C). The optimised IONP concentration of 10 µg/ml (described above) was next used to evaluate uptake rate for IONP for timepoints between 1 and 24 hrs (Figure 4 E, F, respectively). Peak IONP uptake in primary monocytes at 5 hrs of incubation, with uptake plateauing by 6 hrs (Figure 4 F,H), while HUVECs showed time-dependent uptake (Figure 4 E, G). Live video imaging of IONP uptake into both primary monocytes and endothelial cells is demonstrated within the supplementary material (Figure S5).

**Figure 4:**
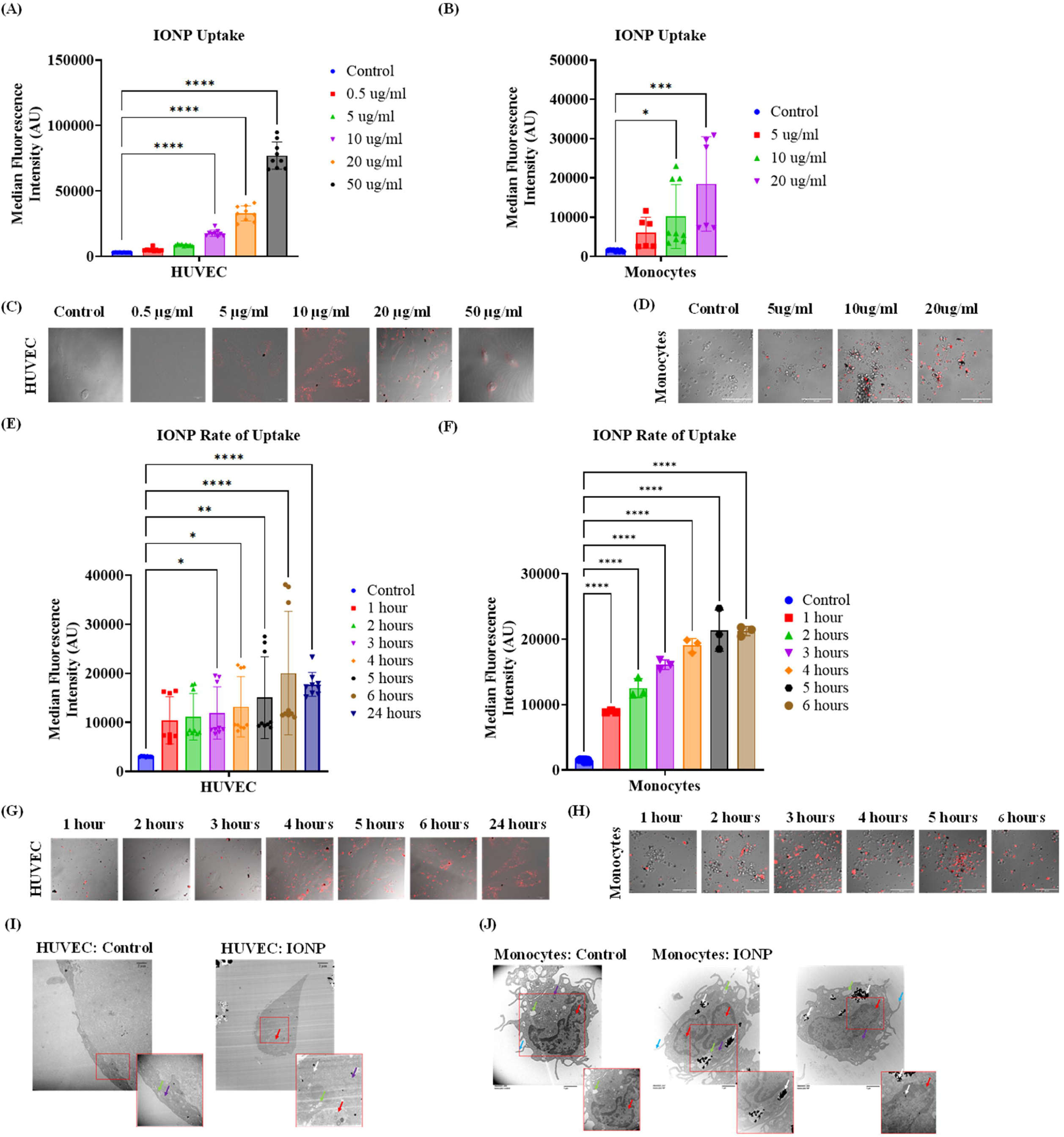
(A, B) Cells were incubated with 0.5, 5, 10, 20 and 50 µg/ml Iron oxide nanoparticle (IONP) uptake was assessed via Flow Cytometry following 24 hrs of incubation in HUVECs and primary monocytes, respectively; (C, D) Fluorescent micrographs showing Iron oxide nanoparticle (IONP) uptake after 24hrs at different concentrations 24 hrs of incubation in HUVECs and primary monocytes, respectively, Scalebar: 50 µM. (E-G, F-H) Cells were incubated with 10 ug/ml of IONP and rate of uptake was assessed from 1-6 hrs and/or 24 hrs via Flow Cytometry and confocal microscopy in HUVECs and primary monocytes, respectively; (I,J) Intracellular location of IONPs following incubation with 10 µg/ml of IONP for 24 hrs visualised by Transmission electron microscopy (TEM) in (I) HUVECs, (J) Primary monocytes. White arrow indicates IONP; red arrow indicates nuclei; blue arrow indicates lipids; purple arrow indicates mitochondria; black arrow indicates RER; green arrow indicates vesicles, Scalebar: 2 µM. Data are represented as mean ± SD, (n=3), statistical comparisons were performed using ANOVA analysis, post-hoc. Statistical significance is denoted as *p<0.05, **p<0.01, ***p<0.005, ****p<0.001

Similarly to ACC cells, TEM images demonstrated cytoplasmic and lysosomal uptake of IONP into endothelial cells (Figure 4I) and primary monocytes (Figure 4J) without uptake into other organelles.

### IONP uptake in ACC cells significantly decreased in the presence of endothelial cells and primary monocytes within a transwell system

At concentrations of 10 µg/ml, IONP uptake into primary monocytes and endothelial cells occurred at a greater rate than into ACC cells. Physiologically, IONP will encounter cellular components of the RES prior to ACC. Therefore, we simulated this environment using a transwell system (Figure 5A) which assessed IONP uptake, in the presence of primary monocytes and an endothelial layer.

**Figure 5:**
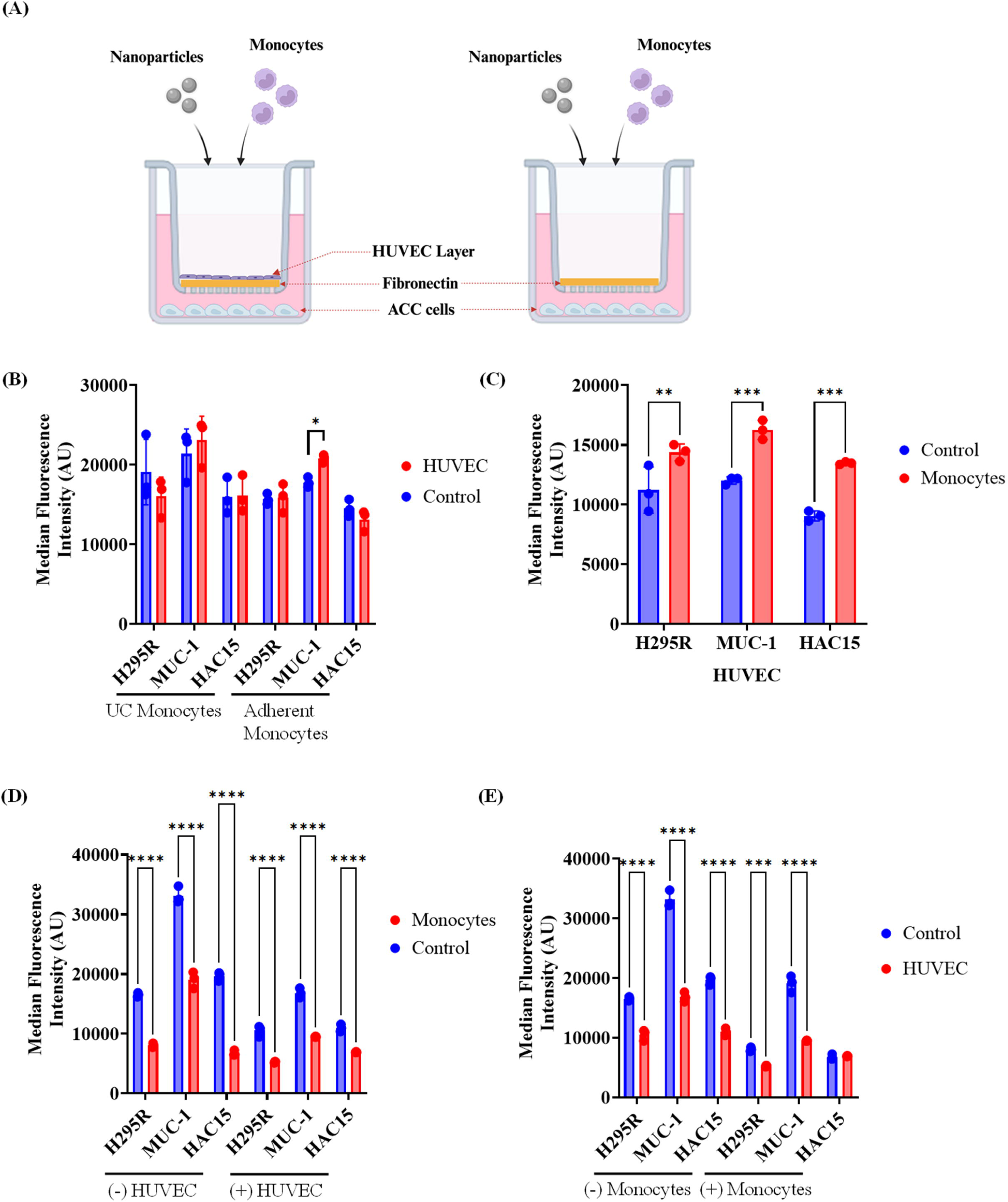
(A) Iron oxide nanoparticle (IONP) uptake by Monocytes and ACC (MUC1, H295R and HAC15) cells across Endothelial and Fibronectin layer using a Transwell migration system; (B) Chamber-specific uptake (MFI) of IONPs into primary monocytes in presence of HUVEC layer and adherent ACC cells in the bottom chamber; (C) Uptake (MFI) of IONPs into HUVECs in presence or absence of Monocytes with adherent ACC cells in the bottom chamber; (D) Uptake (MFI) of IONPs into ACC cells in presence and absence of primary monocytes through Fibronectin layer i.e., (-) HUVEC or endothelial barrier i.e., (+) HUVEC; (E) Uptake (MFI) of IONPs into ACC cells in presence or absence of Monocytes through endothelial barrier.

Primary monocytes which remained in the upper transwell chamber without migrating through the HUVEC layer are denoted as ‘Upper Chamber (UC) monocytes’ and monocytes which migrated through the endothelial barrier and remained adherent in the bottom chamber along with the ACC cells are denoted as ‘Adherent Monocytes’. Uptake of IONPs into the primary monocytes was unaffected by the presence or absence of ACC cells and the endothelial barrier. There was no significant difference in IONP uptake in migrating and non-migrating monocytes (Figure 5B). Uptake of IONPs into HUVECs was significantly higher in the presence of monocytes and ACCs compared to control (Figure 5C). As expected, ACC cell uptake of IONPs for all three cell lines was significantly lower in the individual presence of both monocytes and HUVEC. The combined presence of HUVEC and primary monocytes further reduced the uptake of IONP into all three ACC cell lines. (Figure 5&E). Therefore, while ACC cells avidly take up IONP, this is significantly reduced by non-specific reticuloendothelial cell uptake of IONP.

## Discussion

The current study’s findings demonstrate that iron oxide nanoparticles (IONPs) are avidly taken up by adrenocortical carcinoma (ACC) cells in a time- and concentration-dependent manner, with optimal uptake observed at a concentration of 10 µg/mL. Timelapse imaging demonstrated that the IONP uptake occurred through micropinocytosis. Once taken up, TEM showed that these nanoparticles were located primarily intracytoplasmically and intralysosmally, with no detectable presence in nuclei or mitochondria. IONP at 10 µg/mL did not aggregate in solution and did not demonstrate significant cellular toxicity. Specifically, they did not significantly affect cellular viability, steroidogenesis or metabolic activity, demonstrating their suitability as nanobiocontrast, but also showing that they do not have cytotoxic capacity in the absence of combination therapeutic approaches. Non-specific IONP uptake into monocytes and endothelial cells also occurred and this compromised ACC uptake. Overall, these results highlight the theranostic potential of IONPs in ACC, demonstrating their suitability for diagnostic imaging and also suggesting their suitability as targeted therapeutic agents, as part of combination therapy in future applications.

The uptake of IONPs by ACC cells occurred via macropinocytosis, a process consistent with other studies on nanoparticle internalization in tumour cells. Previous studies have demonstrated the uptake of IONP between 120 and 500nm by through clathrin-mediated endocytosis [19]. In this work, the rate and degree of uptake was different between cell lines and correlated with clathrin expression[20,21]. This may account for the difference in uptake that was observed across the three ACC cell lines. Uptake occurred at a slower rate in the primary tumour cell lines, while the metastatic MUC-1 cell line took up nanoparticles rapidly and actively. Additionally, ACC cells in culture moved towards and engulfed IONP suggesting the additional contribution of a chemotactic mechanism in modulating IONP uptake. The greater avidity of the metastatic cell line for IONPs is encouraging, as targeting metastatic disease will constitute a key theranostic aim for these agents.

The optimised concentration of IONP in this study was 10 µg/mL. Higher concentrations (20 and 50 µg/mL) aggregated within the culture medium due to their inherent magnetic properties and were also toxic to endothelial cells and monocytes [19]. However, at the optimised lower concentration, no cytotoxicity was observed in ACC cell lines. Specifically, the IONP did not affect cellular viability, metabolic activity, proliferation or steroidogenesis. It is interesting that this was matched by intracellular accumulation within the cytoplasm and lysosomes only, without accumulation within the nucleus or mitochondria. Previous studies which have investigated IONP in breast cancer have demonstrated that decreased cellular viability and cell cycle arrest were associated with nuclear accumulation of the nanoparticles[22–24]. There have also been data which have suggested that IONP may demonstrate preferential cytotoxicity in tumour cells [25], where they may more avidly concentrate within the nucleus [24]. We did not see a difference in toxicity in ACC cells versus endothelial cells or monocytes, suggesting that additional strategies are necessary to either accumulate IONP within ACC cells using ligand coating [26,27] or alternatively to use complimentary cytotoxic strategies such as the co-delivery of hyperthermia or cytotoxic chemotherapy [28].

It was interesting that IONP accumulation within ACC cells did not inhibit steroidogenesis. Gonadal accumulation of iron in hereditary haemochromatosis is associated with hypogonadism in certain individuals with poorly controlled disease [29,30]. This has been associated with mitochondrial accumulation of iron, which, again, we did not observe in this study.

Despite the success of IONP uptake in ACC cells, the data presented herein observed significant non-specific uptake by other cell types, including human umbilical vein endothelial cells (HUVECs) and primary human monocytes. The study modelled these cell types in a multicellular, transwell culture system to mimic the endothelial barrier and the expected exposure to the reticuloendothelial system, upon intravenous or systemic administration of IONP in vivo. Under physiological conditions, IONPs will inevitably encounter various cell types in the circulation, and strategies to limit off-target interactions will be crucial to ensure that the nanoparticles reach their intended target in sufficient concentrations [31]. In the current study, uptake of IONP into RES components reduced their uptake into ACC cells. Overall, non-specific IONP uptake presents a considerable challenge for the application of IONPs in the management of cancer and we have shown that this is no different for ACC. For theranostic strategies of IONP to be successful in ACC, it will be crucial to limit off-target uptake by non-ACC cells. Our data demonstrate that not only will non-specific uptake potentially divert IONPs away from an ACC tumour, but additionally, their off-target effects with accumulation may also present off-target cytotoxic effects and hence undesired systemic adverse effects.

It was not the intention of this study to specifically target ACC cells with the bare IONP which we used, but rather to evaluate the uptake and direct toxicity of these nanoparticles to ACC. We know from previous work of members of this group that there is significant RES and liver uptake of these nanoparticles in vivo [32]. For this reason, an in vivo model was not used for evaluation in this study. It is, however, crucial that future studies address strategies which will more specifically target ACC cells using ligand-coating and so-called don’t eat me signals [33–35]. Ligand conjugation, using antibodies or peptides to target specific receptors overexpressed in ACC cells represents one strategy to target IONP to these cells. However, this is challenged by the rarity of ACC and the fact that few phenotyping studies have been carried out which identify specifically overexpressed surface markers on ACC cells. An alternative approach could employ “don’t eat me” signals to prevent phagocytic cells, particularly those in the reticuloendothelial system (RES), from taking up IONPs [34]. One promising candidate for this strategy is the CD47 coating IONP, which inhibits phagocytosis by binding to signal regulatory protein alpha (SIRPα) on macrophages [36]. Coating IONPs with CD47 or similar molecules may help evade uptake by the RES, increasing the likelihood that IONPs will reach the ACC cells. However, the effects of CD47 coating may also inhibit the uptake of IONP into ACC cells, particularly given that micropinocytosis in macrophages uses this pathway [31]. In summary, while IONPs hold promise as theranostic agents for ACC, further research is required to increase their cytotoxicity and to improve specificity for ACC cells. Modifications such as ligand conjugation or RES-evasion strategies will likely be necessary to enhance their therapeutic efficacy and reduce the risk of non-specific uptake. Optimizing these strategies will be a key step in the development of IONPs as a feasible clinical tool for ACC management. Our findings also underscore the complexity of nanoparticle delivery within a biological system and highlight the competitive uptake of IONP by various cellular components, demonstrating the need for careful evaluation of these strategies in vitro before proceeding to in vivo studies.

## Abbreviations

ACC: adrenocortical carcinoma
IONP: iron oxide nanoparticles
PBMCs: peripheral blood mononuclear cells
OCR: oxygen consumption rate
ATP: adenosine triphosphate
FCCP: carbonyl cyanide 4-(trifluoromethoxy) phenylhydrazone
TEM: transmission electron microscopy
PFA: paraformaldehyde
HPLC: high-performance liquid chromatography
DPBS: Dulbecco’s phosphate-buffered saline

## Acknowledgments

The authors would like to thank Coralie Mureau, Catherine Loughney and Mark Webber for their technical guidance. All flow cytometry experiments were performed in the University of Galway Flow Cytometry Core Facility which is supported by funds from University of Galway, Science Foundation Ireland, the Irish Government’s Programme for Research in Third Level Institutions, Cycle 5 and the European Regional Development Fund. Technical and consultative support for flow cytometry experiments was provided by Dr. Shirley Hanley of the University of Galway Flow Cytometry Core Facility.

The authors acknowledge the facilities and scientific and technical assistance of the Centre for Microscopy & Imaging at the University of Galway.

## Author contributions

A.S. and M.C.D. designed the study. S.B. and O.C.Z. designed the nanoparticles. O.C.Z., J.C. and S.V. synthesized and characterized the nanoparticles. A.S. and C.H. performed the cell experiments. P.O provided technical and consultative support for Confocal microscopy. R.S.C, A.S. and C.H. completed data analysis. A.S, R.S.C, C.H, and M.C.D. wrote the manuscript. All authors have contributed to data interpretation and critical revision of the manuscript.

## Data availability

The authors confirm that the data supporting the findings of this study are available within the manuscript and the supplementary materials. Raw data that support findings of this study are available from the corresponding author upon reasonable request.

