## Supplementary material for "Adrenocortical Cancer Cell uptake of Iron Oxide Nanoparticles"

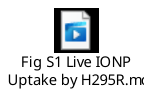

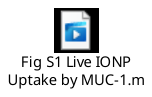

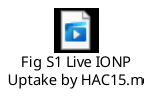

**Figure S1:** Live imaging of iron oxide nanoparticle (IONP) uptake over 24 hours by ACC. ACC cells (MUC1, H295R, HAC15) were incubated with 10 µg/ml of IONP for 24 hours, and uptake was assessed using Confocal Microscopy. An image was taken every 15 minutes over 24 hours. Live imaging videos are in the supplementary section. Scalebar: 50 µm.

Metabolic activity for all cell types was assessed with alamarBlue at day 1, 3 and 7. At day 3, IONP concentration of 50 µg/ml significantly reduced HAC15 metabolic activity (70.96 ±6 % vs 100% control, p<0.05). However, by day 7 metabolic activity of HAC15 at 50 µg/ml showed recovery (Day 3 = 70.96 ±6 % vs Day 7 = 79.4 ± 6 %). Cell metabolic activity remained stable at Day 7 for all cell types (Figure S2A). Seahorse FX24 extracellular analyser was used to assess the effect of the optimum IONP concentration of 10 µg/ml on the respiration of the cells by measuring mitochondrial function using the Mito Stress Test Kit (Figure S2B). Parameters such as spare respiration capacity, maximal respiration, ATP production and non-mitochondrial oxygen consumption were measured for MUC1, H295R and HAC15 cells. 10 µg/ml of IONP did not affect mitochondrial function of the ACC cells (Figure S2C).

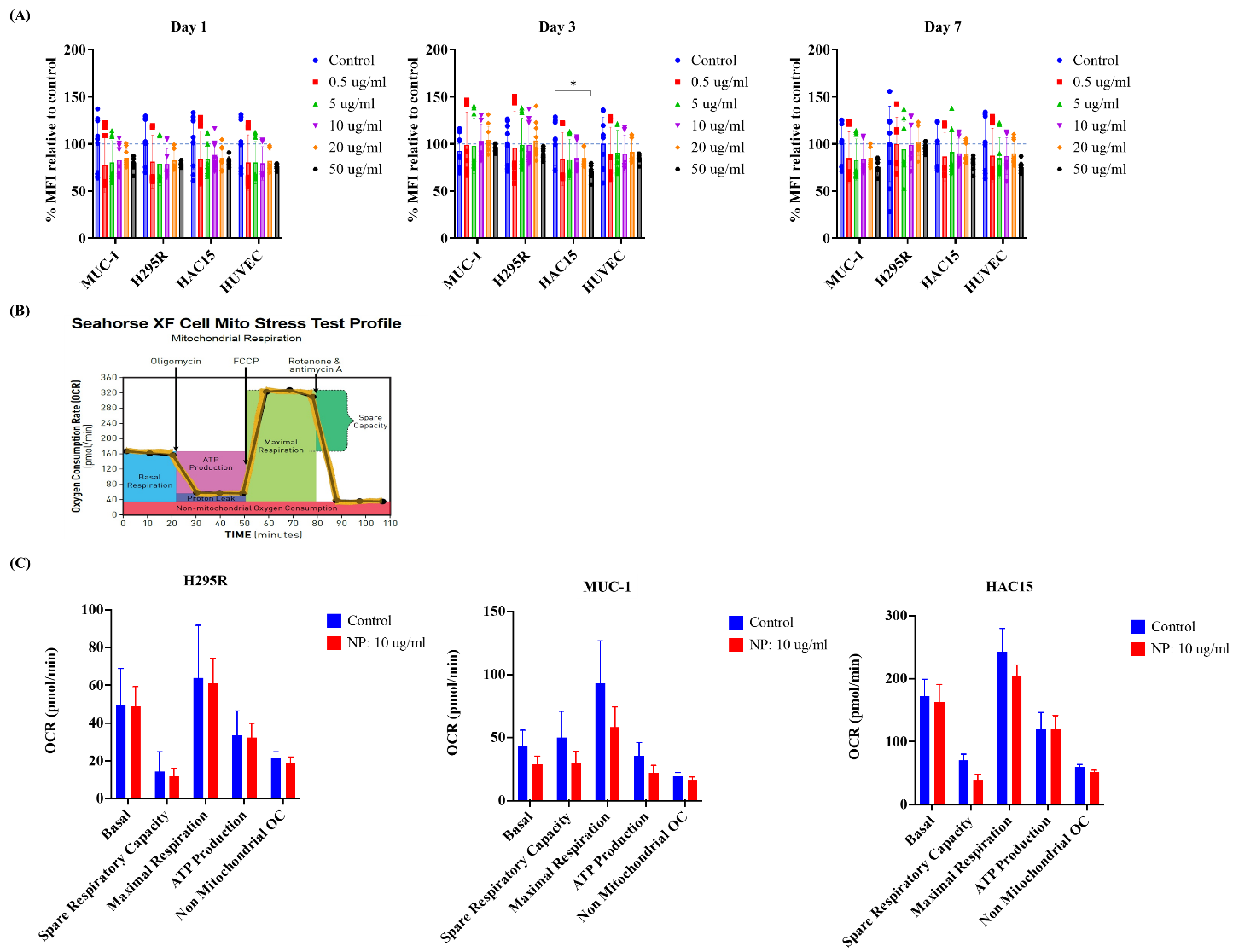

**Figure S2:** (A) Metabolic activity and Seahorse analysis following incubation with 0.5, 5, 10, 20 and 50 µg/ml of IONP for 1, 3 and 7 days using alamarBlue to assess metabolic activity. (B) MUC1, H295R and HAC15 cells were incubated with 10 µg/ml of IONP for 24 hours and mitochondrial respiration was assessed with a Seahorse FX24 extracellular analyser in real time after addition of 2 µM oligomycin, and 0.125, 0.25 and 0.5 µM FCCP. Data are represented as mean ± SD, (n=3), statistical comparisons were performed using ANOVA analysis, post-hoc. Statistical significance is denoted as *p<0.05, **p<0.01, ***p<0.005, ****p<0.001.

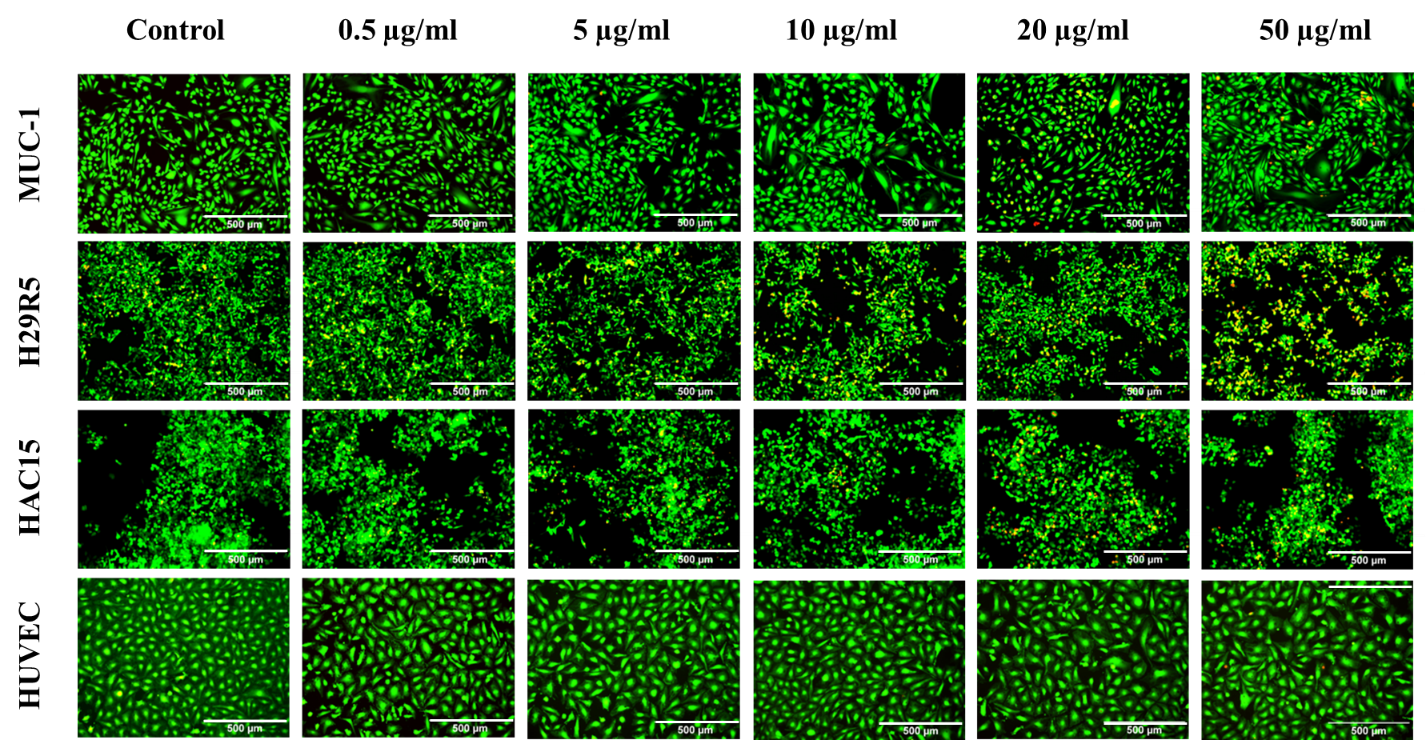

**Figure S3:** Cellular viability following 24 hours incubation. All cell types were incubated with 0.5, 5, 10, 20 and 50 µg/ml of IONP for 24 hours. Viability was assessed with fluorescent microscopy by staining all cell types with calcium AM and ethidium homodimer. Scalebar: 500 µm.

Morphological analysis indicated a change in morphology at 50 µg/ml for HAC15 and HUVEC cells. HAC15 cells showed a more rounded morphology and a reduction in cell spreading (Figure S4A). Ki67 expression analysis was performed to assess the effect of IONP concentration on cell proliferation. Ki-67 expression was not affected in by IONP concentrations of 0.5-50 µg/ml H295R and HAC15. A significant increase in Ki-67 expression was observed by MUC1 following incubation with 10, 20 and 50 µg/ml of IONP. Ki-67 expression was not affected by 10 µg/ml IONP, independent of cell type (Figure S4B,C).

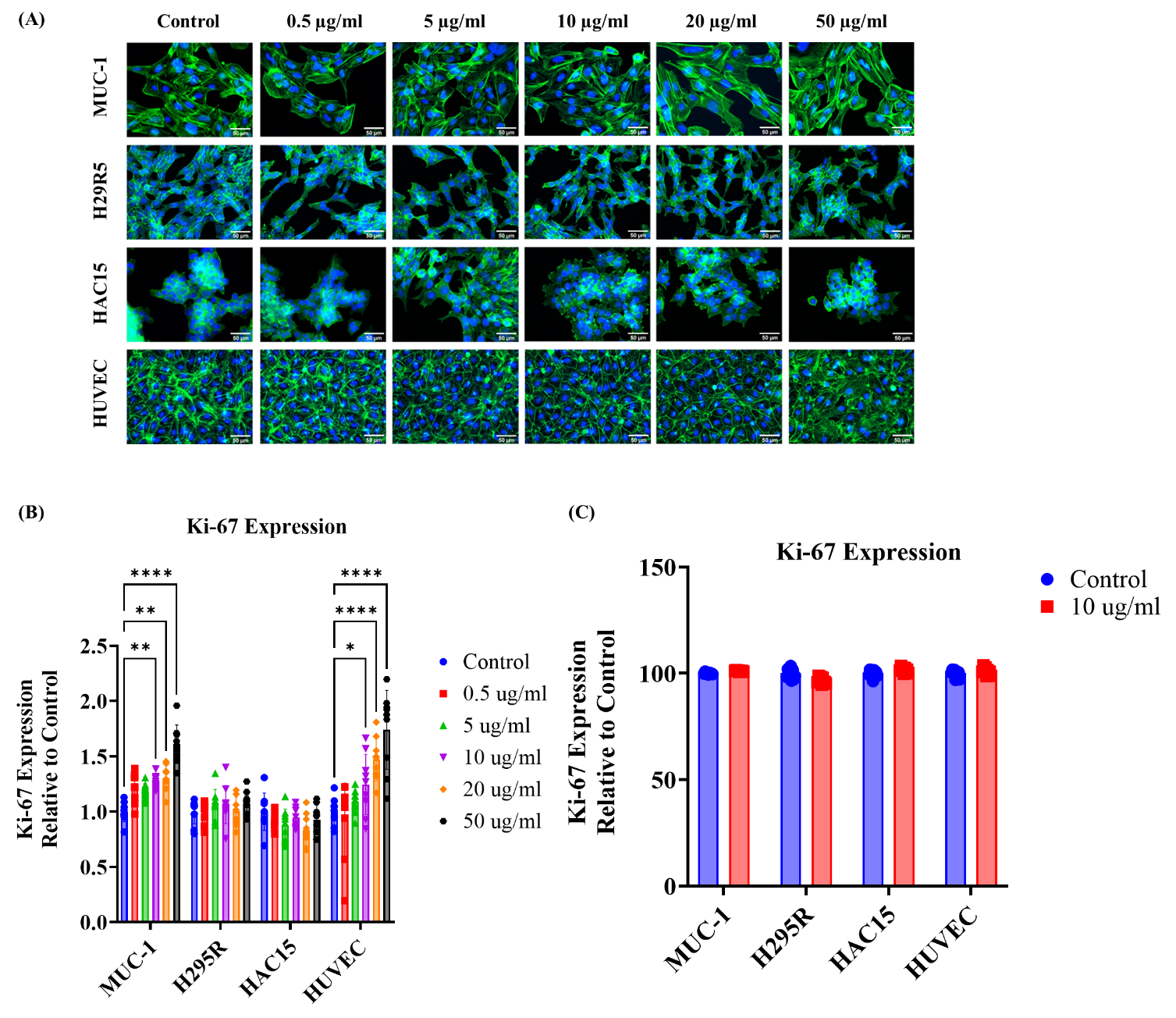

**Figure S4:** (A) Morphological assessment and Ki-67 expression of MUC1, H295R, HAC15 and HUVEC cells following incubation with 0.5, 5, 10, 20 and 50 µg/ml of IONP for 24 hours. Cytoskeleton was stained with FITC-Phalloidin, and nuclei was stained with DAPI. Data are represented as mean ± SD, (n=3), statistical comparisons were performed using ANOVA analysis, post-hoc. Statistical significance is denoted as *p<0.05, **p<0.01, ***p<0.005, ****p<0.001. Scalebar: 50 µM. A change in morphology at 50 µg/ml for HAC15 and HUVEC cells was demonstrated. HAC15 cells showed a more rounded morphology and a reduction in cell spreading; (B) Cell proliferation was assessed by Ki-67 staining following 24hrs, represented as MFI (AU) vs control; Ki-67 expression was not affected in by IONP concentrations of 0.5-50 µg/ml in H295R and HAC15. A significant increase in Ki-67 expression was observed in MUC1 following incubation with 10, 20 and 50 µg/ml of IONP. (C) Cell proliferation was assessed by Ki-67 staining following incubation with 10 µg/ml of IONP for 7 days, represented as MFI (AU) vs control. Ki-67 expression was not affected by 10 µg/ml IONP, independent of cell type.

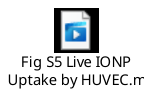

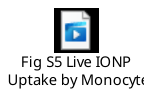

**Figure S5:** Live imaging of IONP uptake over 24 hours by Monocytes and HUVEC cells**.** Monocytes and HUVEC cells were incubated with 10 µg/ml of IONP and uptake was assessed using Confocal Microscopy. An image was taken every 15 minutes over 24 hours. Live imaging videos are in the supplementary section. Scalebar: 50 µm.
